# Sex-specific impact of prenatal androgens on intrinsic functional connectivity between social brain default mode subsystems

**DOI:** 10.1101/253310

**Authors:** Michael V. Lombardo, Bonnie Auyeung, Tiziano Pramparo, Angélique Quartier, Jérémie Courraud, Rosemary J. Holt, Jack Waldman, Amber N. V. Ruigrok, Natasha Mooney, Meng-Chuan Lai, Prantik Kundu, Edward T. Bullmore, Jean-Louis Mandel, Amélie Piton, Simon Baron-Cohen

## Abstract

Many early-onset neurodevelopmental conditions such as autism affect males more frequently than females and affect corresponding domains such as social cognition, social-communication, language, emotion, and reward. Testosterone is well-known for its role as a sex-related biological mechanism and affects these conditions and domains of functioning. Developmentally, testosterone may sex-differentially impact early fetal brain development by influencing early neuronal development and synaptic mechanisms behind cortical circuit formation, particularly for circuits that later develop specialized roles in such cognitive domains. Here we find that variation in fetal testosterone (FT) exerts sex-specific effects on later adolescent functional connectivity between social brain default mode network (DMN) subsystems. Increased FT is associated with dampening of functional connectivity between DMN subsystems in adolescent males, but has no effect in females. To isolate specific prenatal neurobiological mechanisms behind this effect, we examined changes in gene expression identified following a treatment with a potent androgen, dihydrotestosterone (DHT) in an in-vitro model of human neural stem cell (hNSC). We previously showed that DHT-dysregulates genes enriched with known syndromic causes for autism and intellectual disability. DHT dysregulates genes in hNSCs involved in early neurodevelopmental processes such as neurogenesis, cell differentiation, regionalization, and pattern specification. A significant number of these DHT-dysregulated genes shows spatial expression patterns in the adult brain that highly correspond to the spatial layout of the cortical midline DMN subsystem. These DMN-related and DHT-affected genes (e.g., *MEF2C*) are involved in a number of synaptic processes, many of which impact excitation/inhibition imbalance. Focusing on *MEF2C*, we find replicable upregulation of expression after DHT treatment as well as dysregulated expression in induced pluripotent stem cells and neurons of individuals with autism. This work highlights sex-specific prenatal androgen influence on social brain DMN circuitry and autism-related mechanisms and suggests that such influence may impact early neurodevelopmental processes (e.g., neurogenesis, cell differentiation) and later developing synaptic processes.

---

It has long been known that events occurring during prenatal development can have long-lasting programming impact on susceptibility for medical conditions that emerge later in life^1-3^. Emerging work has shown that events starting during prenatal development can have long-lasting impact potentially leading to increased risk for atypical neurodevelopmental phenotypes. For instance, neurodevelopmental conditions such as autism have been linked to multiple types of biological processes that stem from very early periods of prenatal development (e.g., ^4-10^). Therefore, it is becoming increasingly clear that fetal brain development is a critical developmental window of importance for understanding factors that increase the likelihood of neurodevelopmental conditions.

Several early-onset neurodevelopmental conditions (e.g., autism, intellectual disability, ADHD, developmental language disorders, conduct disorder) are well known to have a sex-biased prevalence rate^11^. For example, the latest estimates for autism suggest that 3 males are affected for every 1 female^12^. Several theories have been put forward to explain the sex ratio imbalance in early-onset neurodevelopmental conditions ^13^. One prominent theory suggests that there are factors inherent in females that act to reduce the likelihood of atypical neurodevelopment^14^. For example, a higher burden of large effect mutations are present in females compared to males with autism, suggesting that protective mechanisms in females may raise the threshold for deleterious impact of such mutations^15^. In contrast to female-specific factors reducing likelihood of autism, there may also be important male-specific mechanisms for increasing likelihood of autism. We have theorized that sex steroid hormones such as testosterone may have fetal programming impact on later brain development in ways relevant to autism^16^. Recent large studies have confirmed that maternal polycystic ovary syndrome, a syndrome associated with elevated androgen levels, increases the odds for both autism and ADHD in offspring^17, 18^. We also recently confirmed elevation of a latent steroidogenic factor spread across multiple steroid hormones including testosterone in the amniotic fluid of clinically diagnosed males with autism^7^. Extending into the non-clinical population, continuous variation in how much fetal testosterone (FT) an individual is exposed to in the womb is associated with behavioral variation in autistic traits^19, 20^. FT and later postnatal testosterone variation also affects specific behavioral and cognitive domains such as social cognition, language, emotion, and reward processes, which are implicated in autism and other male-biased neurodevelopmental conditions^21, 22^. Thus, it may be that multiple sex-specific factors could be at work to both increase likelihood in males and decrease likelihood in females for developing early-onset neurodevelopmental conditions. Further work is needed to tease apart how these mechanisms may operate similarly or differently in males and females. In addition, translational work is needed to examine how such mechanisms directly affect biological processes supporting both typical and atypical human brain development.

In this study, we first examine the question of whether FT longitudinally exerts sex-specific influence over intrinsic functional organization of specific circuits involved in social, language, and affective functions in humans. To test this question we examined a unique cohort of individuals in whom we have measured concentration levels of testosterone directly from amniotic fluid during a midgestational window of pregnancy where fetuses are sexually differentiating and where surges in sex steroid levels may have maximal impact on early brain development^23, 24^. These individuals are now in their adolescent years and we examined how variability in FT is associated with patterns of intrinsic functional connectivity as measured with resting state fMRI (rsfMRI). Given known links between FT and behavioral and cognitive domains of social cognition, social-communication, language, emotion, and reward^21, 22^, we examined specific neural circuits known for their roles in these domains. We predicted that FT would act as a male-specific risk mechanism to influence connectivity towards atypicality and that such male-specific influence would not be observed in females. To further understand the underlying biology behind FT influence on neural circuitry, we also examined androgen-related changes in gene expression in human neural stem cells (hNSC) recently reported^25^ and the overlap between these androgen-sensitive genes and genes spatially expressed in similar patterns across cortex as rsfMRI-defined macroscale networks.

## Methods

### Participants

This study was approved by the Essex 1 National Research Ethics committee. Parents gave informed consent for their child to participate and each adolescent also gave assent to participate. Participants were 64 adolescents (32 males, 32 females; male age mean = 15.42 years, standard deviation = 0.92 years; female age mean = 15.55 years, standard deviation = 1.06 years; age range = 13.22-17.18 years) sampled from a larger cohort of individuals whose mothers underwent amniocentesis during pregnancy for clinical reasons (i.e. screening for chromosomal abnormalities). At amniocentesis, none of the individuals screened positive for any chromosomal abnormalities and were thus considered typically developing. At the time of scanning, none of the participants self- or parent-reported any kind of neurological or psychiatric diagnosis. After assays of current testosterone levels were completed, we found that the assay did not result in useable data for 4 males and 2 females, and thus these individuals were excluded from further analyses requiring intact FT and current testosterone data.

### Fetal testosterone collection and measurement

FT was measured from amniotic fluid samples collected between 13 and 20 weeks of gestation via radioimmunoassay. This period is within the 8–24 week window that is hypothesized to be critical for human sexual differentiation^23^. Amniotic fluid was extracted with diethyl ether, which was evaporated to dryness at room temperature and the extracted material redissolved in an assay buffer. Testosterone was assayed by the Count-a-Coat method (Diagnostic Product), which uses an antibody to testosterone coated onto propylene tubes and a 125I- labeled testosterone analog. The detection limit of the assay using the ether-extraction method is 0.05 nmol/L. The coefficient of variation (CV) for between-batch imprecision is 19% at a concentration of 0.8 nmol/L and 9.5% at a concentration of 7.3 nmol/L. The CVs for within- batch imprecision are 15% at a concentration of 0.3 nmol/L and 5.9% at a concentration of 2.5 nmol/L. This method measures total extractable testosterone.

### Current testosterone collection and measurement

Current testosterone during adolescence was measured from passive saliva samples using a commercial competitive ELISA (Salimetrics Ltd, US) at the Biomarker Analysis Laboratory at Anglia Ruskin University. All pipetting used a Tecan Evo liquid handler. 25μl of saliva samples, standard and controls were pipetted into the appropriate wells of the ELISA plates, 150μl of testosterone conjugated to horseradish peroxidase (HRP) enzymes was added to all wells and the plate shaken for 5 minutes and then incubated for 55 minutes at room temperature. The ELISA plates were washed four times in Salimetrics wash buffer using a Tecan Hydroflex before the addition of 200μl of enzyme substrate (TMB), the plates shaken for 5 minutes and colour allowed to develop for an additional 25 minutes before the reaction was stopped by the addition of 50μl of Salimetrics stop solution. The resulting optical density in the wells was read at 450nm with a secondary filter correction at 620nm. The sensitivity of this assay is 1pg/ml. The CVs for within-batch imprecision were 4.53% for the high controls and 14.88% for the low controls.

### fMRI Data Acquisition

All MRI scanning took place on a 3T Siemens Tim Trio MRI scanner at the Wolfson Brain Imaging Centre in Cambridge, UK. Functional imaging data were acquired with a multi-echo EPI sequence with online reconstruction (repetition time (TR), 2000 ms; field of view (FOV), 240 mm; 28 oblique slices, descending alternating slice acquisition, slice thickness 3.8 mm; TE = 13, 31, and 48 ms, GRAPPA acceleration factor 2, BW = 2368 Hz/pixel, flip angle, 90°, voxel size 3.8mm isotropic). Resting state data were collected using a 10 minute ‘eyes-open’ run (i.e. 300 volumes), where participants were asked to stare at a central fixation cross and to not fall asleep. Anatomical images were acquired using a T1-weighted magnetization prepared rapid gradient echo (MPRAGE) sequence for warping purposes (TR, 2300 ms; TI, 900 ms; TE, 2.98 ms; flip angle, 9°, matrix 256 × 256 × 256, field-of-view 25.6 cm).

### fMRI Preprocessing

Data were processed by ME-ICA using the tool meica.py as distributed in the AFNI neuroimaging suite (v2.5), which implemented both basic fMRI image preprocessing and decomposition-based denoising. For the processing of each subject, first the anatomical image was skull-stripped and then warped nonlinearly to the MNI anatomical template using AFNI *3dQWarp*. The warp field was saved for later application to functional data. For each functional dataset, the first TE dataset was used to compute parameters of motion correction and anatomical-functional coregistration, and the first volume after equilibration was used as the base EPI image. Matrices for de-obliquing and six-parameter rigid body motion correction were computed. Then, 12-parameter affine anatomical-functional coregistration was computed using the local Pearson correlation (LPC) cost function, using the gray matter segment of the EPI base image computed with AFNI *3dSeg* as the LPC weight mask. Matrices for de-obliquing, motion correction, and anatomical-functional coregistration were combined with the standard space non-linear warp field to create a single warp for functional data. The dataset of each TE was then slice-time corrected and spatially aligned through application of the alignment matrix, and the total nonlinear warp was applied to the dataset of each TE. No time series filtering was applied in the preprocessing phase. Data were analyzed with no spatial smoothing. ME-ICA denoising was used to identify and remove non-BOLD signal fluctuation^26-28^.

### Group Independent Components Analysis and Dual Regression

To assess large-scale intrinsic functional organization of the brain we utilized the unsupervised data-driven method of independent component analysis (ICA) to conduct a group-ICA, and then utilized dual regression to back-project spatial maps and individual time series for each component and subject. Both group-ICA and dual regression was implemented with FSL’s MELODIC and Dual Regression tools (www.fmrib.ox.ac.uk/fsl). For group-ICA, we constrained the dimensionality estimate to 30, as in most cases with low-dimensional ICA, the number of meaningful components can be anywhere from 10-30^29^. From these components we manually selected the components that best represented networks typically involved in emotion (amygdala), reward (ventral striatum), language (superior temporal gyrus, inferior frontal gyrus, insula), and social cognition (default mode network) functions.

### Between-Component Connectivity Analysis

Time courses for each subject and each component were used to model between-component connectivity. This was achieved by constructing a partial correlation matrix for all 6 components using Tikhonov-regularization (i.e. ridge regression, rho = 1) as implemented within the *nets_netmats.m* function in the FSLNets MATLAB toolbox (https://fsl.fmrib.ox.ac.uk/fsl/fslwiki/FSLNets). The aim of utilizing partial correlations was to estimate direct connection strengths in a more accurate manner than can be achieved with full correlations, which allow more for indirect connections to influence connectivity strength^29-31^. Partial correlations were then converted into Z-statistics using Fisher’s transformation for further statistical analyses. The lower diagonal of each subject’s partial correlation matrix was extracted for a total of 15 separate component-pair comparisons. For each of these 15 comparisons we ran robust regression (to be insensitive to outliers)^32^ (https://github.com/canlab/RobustToolbox) to examine correlations between connectivity and FT variation. This analysis was conducted twice - once with FT only and again with FT and controlling for variability in current levels of testosterone. These correlations were computed separately for males and females. Finally, we computed Z-statistics and p-values for difference between male and female correlations utilizing the *paired.r* function in the *psych* R library. False positive control was achieved with FDR q<0.05, implemented with the *p.adjust* function in R.

### Androgen Influence on Gene Expression in a Human Neural Stem Cell Model

To gain insight into the impact of androgens have on gene expression in embryonic development, we examined data from a recent RNA-seq experiment whereby human neural stem cells (hNSC) derived from embryonic stem cells were treated with 100 nM of a potent androgen, dihydrotestosterone (DHT) or a control treatment (Dimethyl sulfoxide; DMSO)^25^ (GEO accession number: GSE86457). The cell line used was line SA001 from a male donor, obtained from Cellartis (Goteborg, Sweden). The work was supervised by the French Bioethics Agency (Agreement number NOR AFSB 12002 43S). hNSCs were derived as described in Boissart et al.,^33^. Read counts obtained from RNA sequencing of three batches (biological replicates) of treated hNSCs derived from SA001 (DMSO (n = 4;3;3) vs. DHT 100nM (n = 3;3;3)) were analyzed, scaling them by library size using the TMM method, implemented with the calcNormFactors function in the edgeR library^34^. Low expressing genes were removed if there was not 2 or more samples with more than 100 reads while the previous analysis^25^ filtered out genes with less than 100 reads normalized and divided by gene length in kb for each condition and below the 80th percentile in one of the condition studied (DMSO or DHT 100nM 24h). This filtering left a total of 13,284 genes for further downstream analysis. Batch effects were removed using the *ComBat* function within the *sva* R library. The software utilized in the prior paper^25^ for DE analysis was DESeq2^35^.

This study used the *voom* function from the *limma* library in R to estimate precision weights for linear modeling of differential expression (DE) that will account for mean-variance trends^36^. DE analysis in *limma* allows for utilization of the *voom* function to estimate precision weights that account for mean-variance trends and which are incorporated directly into linear DE models. Another unique aspect of DE analysis within *limma* is the incorporation of an empirical Bayes procedure for borrowing information between genes in estimating variance. Genes were identified as DE if they survived Storey FDR correction at q<0.05^37^, instead of the Benjamini and Hochberg FDR method^38^ previously used by Quartier et al.,^25^.

### Isolating DMN-relevant genes based on spatial expression similarity to rsfMRI components

We next wanted to isolate genes with high-relevance to our specific large-scale neural circuits identified by rsfMRI ICA analysis. To achieve this aim, we used the gene expression decoding functionality within Neurosynth and NeuroVault^39^ to identify genes whose spatial expression patterns are consistently similar across subjects to our rsfMRI IC maps. This decoding analysis utilizes the 6 donor brains from the Allen Institute Human Brain Gene Expression atlas^40^, ^41^. The analysis first utilizes a linear model to compute similarity between the observed rsfMRI IC map and spatial patterns of gene expression for each of the six brains in the Allen Institute dataset. The slopes of these subject-specific linear models encode how similar each gene’s spatial expression pattern is with our rsfMRI IC maps. These slopes were then subjected to a one-sample t-test to identify genes whose spatial expression patterns are consistently of high similarity across the donor brains to the rsfMRI IC maps we input. The resulting list of genes was then thresholded for multiple comparisons and only the genes surviving FDR q<0.05 and with positive t-statistic values were considered.

### Gene Set Enrichment Analyses

Analyses examining enrichment (i.e. overlap) between two lists of genes, was implemented using the *sum(dhyper())* function in R. For enrichments with Neurosynth Gene Expression Decoding analyses, the background set size for all enrichment analyses was set to the total number of genes considered for Neurosynth Gene Expression Decoding analyses (i.e. 20,787). We also examined enrichment with autism-associated genes listed under SFARI Gene Scoring categories (lists downloaded on 10/10/2017). The background set size for this analysis was set to the total number of genes included in the DE analysis (13,284).

All enrichment analyses using Gene Ontology (GO) database was implemented with AmiGO 2 (http://amigo.geneontology.org/amigo). Here we used a custom background list, which was the total number of genes included in the DE analysis (i.e. 13,284 genes). Only GO terms surviving Bonferroni correction were used.

### Developmental trajectory of MEF2C expression

RNA-seq data from the Allen Institute BrainSpan Human Brain Gene Expression atlas was utilized for this analysis. We examined medial prefrontal cortex (MFC) in BrainSpan as it is the most prominent region from the IC01 functional connectivity map (see Results). We tested hypotheses about whether there is developmental upregulation of *MEF2C* and enhanced variability in prenatal versus postnatal development. To test this, we used a permutation test (100,000 permutations) whereby on each permutation we randomized prenatal or postnatal labels and then re-calculated the mean difference or difference in standard deviation in expression between prenatal and postnatal (after birth) periods. We then compared the observed mean or standard deviation difference statistics to their null distributions to compute the p-value for each comparison.

### Follow-up experiment of DHT-dysregulation of MEF2C expression using qPCR

Three lines of hNSCs cells (two derived from iPSC lines reprogrammed from male fibroblasts GM01869 and GM04603 and one derived from blood of an anonymous female donor, PB12) were treated by DMSO or DHT 100nM during 24h, in quadruplicates, as previously described in Quartier et al.,^25^. RNA was extracted using RNeasy minikit (Qiagen, Valencia, CA, USA) including a DNase I treatment. Total RNA (500 ng) was reverse transcribed into cDNA using random hexamers and SuperScript II reverse transcriptase according to manufacturer’s recommendation (Invitrogen, Carlsbad, CA, USA). Real-time PCR quantification (qPCR) was performed on cDNA on LightCycler 480 II (Roche) using the QuantiTect SYBR Green PCR Master Mix (Qiagen) and primers specific to *MEF2C* (MEF2C_RT_F: 5’- ATCGACCTCCAAGTGCAGGTAACA-3’ and MEF2C_RT_R: 5’- AGACCTGGTGAGTTTCGGGGATT-3’). All qPCR reactions were performed in triplicate. Reaction specificity was controlled by post-amplification melting curve analysis. The relative expression of gene-of-interest *vs.* two references genes (*GAPDH* and *YWHAZ*) was calculated using the 2-(ΔΔCt) method. To test for DHT upregulation of *MEF2C* expression we utilized a linear mixed-effect model (*lme* function within the *nlme* R library) with condition and sex as fixed effects and cell line and replicate as crossed random effects. To quantify evidence of replication of the original *MEF2C* result in RNA-seq data, we computed a replication Bayes Factor^42^. Replication Bayes Factors greater than 10 indicate strong evidence for replication.

### MEF2C expression in an iPSC model of autism

*MEF2C* expression was examined from the RNA-seq data from Marchetto et al.,^9^. This dataset comprises expression measured from induced pluripotent stem cells (iPSC), neural progenitor cells, and neurons grown from fibroblasts of typically developing controls or patients with autism. Analyses were specifically focused on *MEF2C* expression and used a linear mixed-effect model ANOVA (*lme* function from the *nlme* R library) modeling diagnosis, RIN, cell type, and diagnosis^*^cell type interaction as fixed effects and subject identifier as a random effect. This ANOVA was followed-up by specific tests (*lm* function in R) of between-group difference within each cell type, covarying for RIN.

## Results

### Sex-differential FT influence on connectivity between social brain default mode subsystems

Confirming our overall hypothesis that FT exerts sex-specific influence over connectivity between neural circuits underpinning functions affected in male-biased neurodevelopmental conditions, we find only one between-component connection that is differentially related to FT in males versus females. This between-component connection comprises anterior and primarily cortical midline (IC01) versus posterior (IC09) components of the default mode network (DMN) (Fig 1A-B). On average, connectivity between these components is robustly non-zero indicating strong normative relationships between these two integral parts of the DMN. However, as FT increases, connectivity between IC01 and IC09 decreases, specifically for males but not females (male r = -0.69, female r = 0.02, z = 3.35, p = 7.88e-4) (Fig 1C). In other words, within males specifically, increasing FT has an effect of dampening connectivity between these DMN subsystems. This effect was robust after taking current levels of testosterone into account (z = 3.12, p = 0.001), indicating that this effect is not explained simply by variability in current levels of testosterone during adolescence.

**Figure 1:**
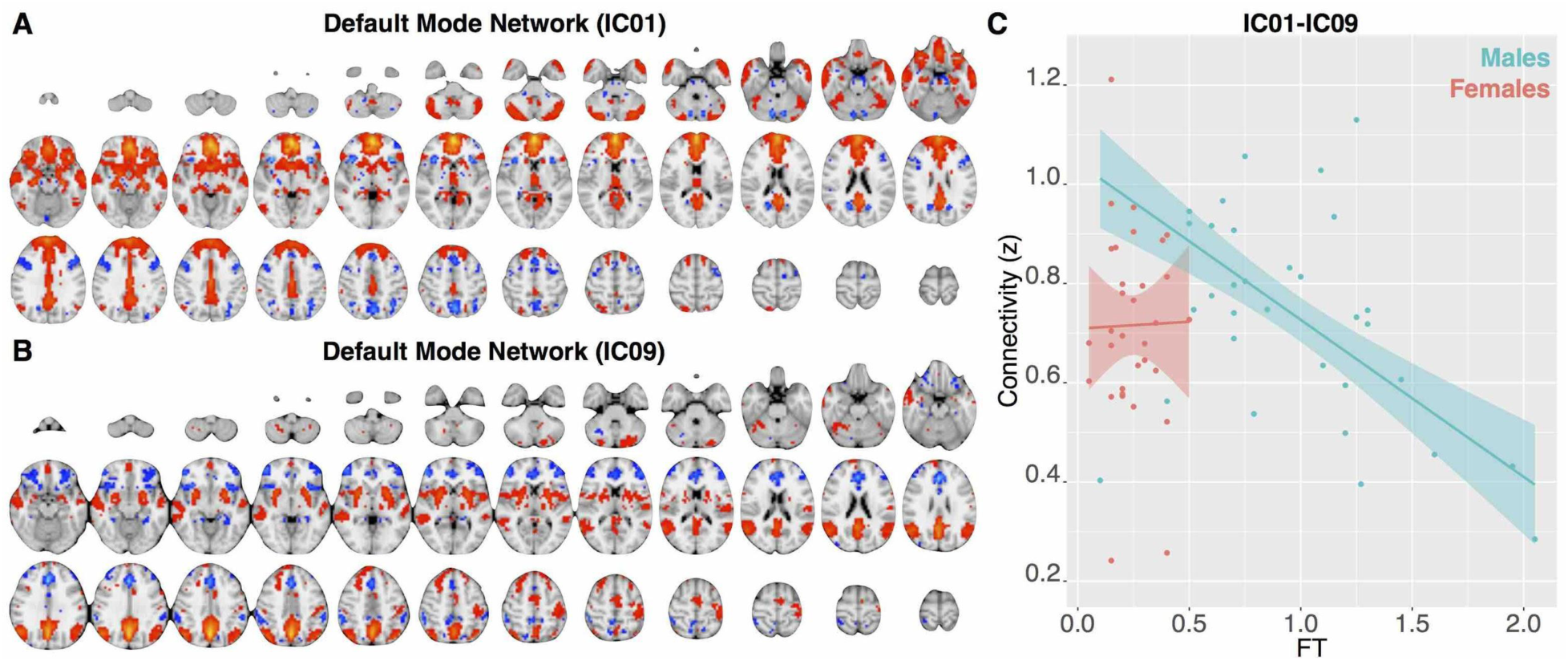
Sex-differential relationship between FT and DMN subsystem connectivity. Panels A and B show axial montages of the two DMN components (IC01, IC09). Panel C shows a scatterplot of the relationship between FT and IC01-IC09 connectivity.

### Genes influenced by androgens and relevant to DMN circuits

We next asked the question of how androgens may impact gene expression in early prenatal development - specifically DMN circuitry. To model these early stages of development, we re-analysed previously published^25^ transcriptomic data obtained from human neural stem cell model (hNSC) treated with a potent androgen, dihydrotestosterone (DHT, 100nM). Differential expression analysis identified 460 genes upregulated and 221 genes downregulated by DHT. Upregulated genes are enriched in numerous processes spanning angiogenesis, blood vessel morphogenesis and development, enzyme linked receptor protein signaling, cell surface receptor signaling, signal transduction, cell morphogenesis, neuron development, and cell differentiation, amongst many others. Downregulated genes are enriched in cardiac chamber morphogenesis, regionalization, pattern specification process, neuron differentiation, neurogenesis, amongst many others (Supplementary Figure 1A-B). We previously reported enrichments with many autism-associated genes which are also known as syndromic causes of autism^25^. Although our DE analyses differed from the past paper^25^, the current enrichment analyses largely confirm this prior finding. Many genes listed in the ‘Syndromic’ category of SFARI Gene were also DHT-dysregulated (e.g., *HI1*, *ASXL3*, *CHD2*, *DHCR7*, *HCN1*, *MEF2C*, *PAX6*, *PRODH*, *PTEN*, and *SCN1A*). Other important autism-associated genes were also DHT-dysregulated such as *NLGN4X*, *NRXN3*, *FOXP1*, and *SCN9A*. For more on this enrichment analysis between DHT-dysregulated genes and autism-associated genes for this analysis, see the Supplementary Results and Supplementary Figure 1C.

We then tested the critical question regarding whether any genes influenced by DHT in hNSCs are relevant for developing cortical networks such as the DMN, which are influenced by FT in a sex-specific manner. We used Neurosynth Gene Expression Decoding analyses to isolate spatial gene expression patterns that are similar to the DMN components identified by rsfMRI. While only 4 genes pass at FDR q<0.05 for the IC09 component, 2,444 genes pass at the same FDR threshold for the IC01 component. This IC01 gene set was significantly enriched in genes that are differentially expressed by DHT in hNSCs (OR = 1.88, p = 0.000002). This overlapping gene set was highly enriched in a number of synaptic processes (Figure 2A). Several examples of such genes are shown in Figure 2B-F - particularly myocyte enhancer factor 2C (*MEF2C*), synaptotagmin 17 (*SYT17*), neurabin-1 (*PPP1R9A*), neuronal pentraxin 2 (*NPTX2*), glutamate receptor 1 (*GRIA1*), and glutamate receptor, ionotropic, kainate 2 (*GRIK2*), sodium channel, voltage gated, type III alpha subunit (*SCN3A*), and sodium channel, voltage gated, type IX alpha subunit (*SCN9A*).

**Figure 2:**
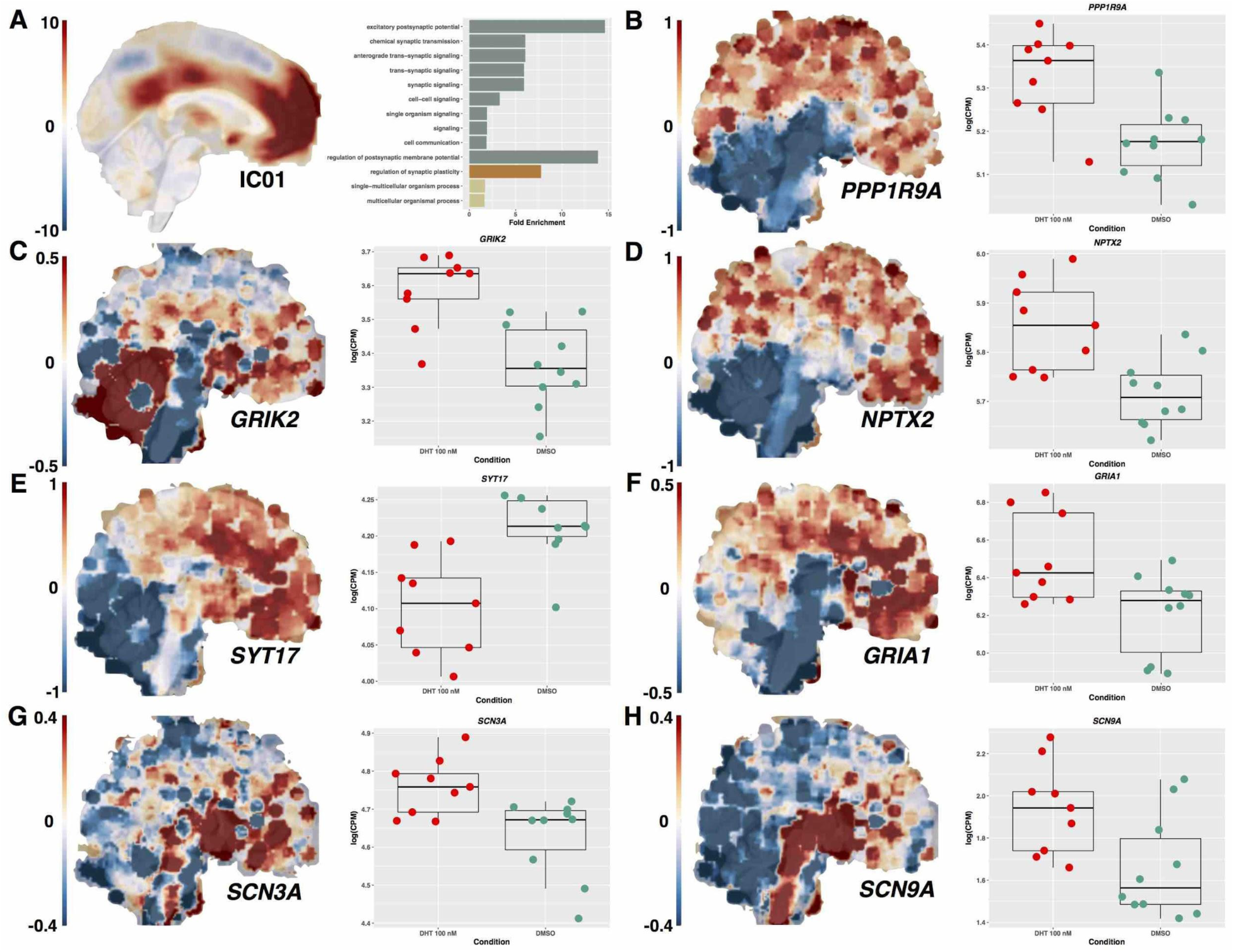
Synaptic enrichments of DHT-dysregulated genes and genes with high levels of spatially expression similarity to rsfMRI DMN IC01 map (A) and plots of specific genes contributing to these enrichments (B-H). Whole-brain maps showing expression for each gene are composite maps averaging across all donors. These composite maps are shown for visualization purposes only. They are not meant to reflect directly the hierarchical statistical testing as implemented with Neurosynth Gene Expression Decoding.

Of the genes that are both differentially expressed by DHT and spatially highly expressed in a pattern associated with the DMN, we followed up one gene of particular interest - *MEF2C* (Figure 3A). *MEF2C* was among the genes from SFARI Gene noted as a syndromic cause of autism and is differentially expressed here by DHT. *MEF2C* is also known to be a downstream target of the androgen receptor^43^. As a transcription factor itself, *MEF2C* is also known to differentially target other genes as a function of sex and the degree of such sex-differential targeting explains a substantial amount of variance in sex-differential expression^44^. Confirming DHT-upregulation of *MEF2C* expression we ran a follow-up experiment using qPCR and found that indeed the upregulation of *MEF2C* by DHT in the RNA-seq data is replicated using qPCR on three additional independent cell lines and (*t* = 3.73*, p* = 0.003*, replication Bayes Factor* = 164) (Figure 3B-C).

We went further in subsequent analyses to identify *MEF2C*’s trajectory of expression in medial prefrontal cortex throughout prenatal and postnatal periods. Using the Allen Institute BrainSpan Developmental Gene Expression atlas we indeed find that fetal development is a prominent period where *MEF2C* is upregulated in medial prefrontal cortex (p = 0.01). There is also enhanced variability in prenatal *MEF2C* expression compared to postnatal expression (p = 0.02) - an effect likely due to marked change from first to second and third trimesters of gestation. *MEF2C* expression increases substantially from first to second trimester of gestation and continues high levels of expression until around 2 years of age, whereby expression tapers off and becomes stable throughout the rest of the lifespan (Fig 3D).

Finally, we tested whether the directionality of *MEF2C* dysregulated gene expression in autism induced pluriopotent stem cells (iPSC) is congruent with the directionality of DHT-influence on *MEF2C* expression. To achieve this aim, we re-examined a recent RNA-seq dataset which examined iPSC, neural progenitor cells, and neurons grown from fibroblasts of male cases of autism^9^. An ANOVA examining all cell types identified a Diagnosis^*^Cell Type interaction (*F* = 6.89, *p* = 0.0047). Congruent with the directionality of DHT-upregulation, *MEF2C* is also upregulated in autism iPSCs (*t* = 3.68, *p* = 0.0035, *Cohen’s d* = 1.42). In contrast, no dysregulation is observed in neural progenitor cells (*t* = 0.12, *p* = 0.90, *Cohen’s d* = 0.11), while in neurons there is a trend for reduced expression in autism (*t* = 2.19, *p* = 0.052, *Cohen’s d* = 1.31) (Fig 3E).

**Figure 3:**
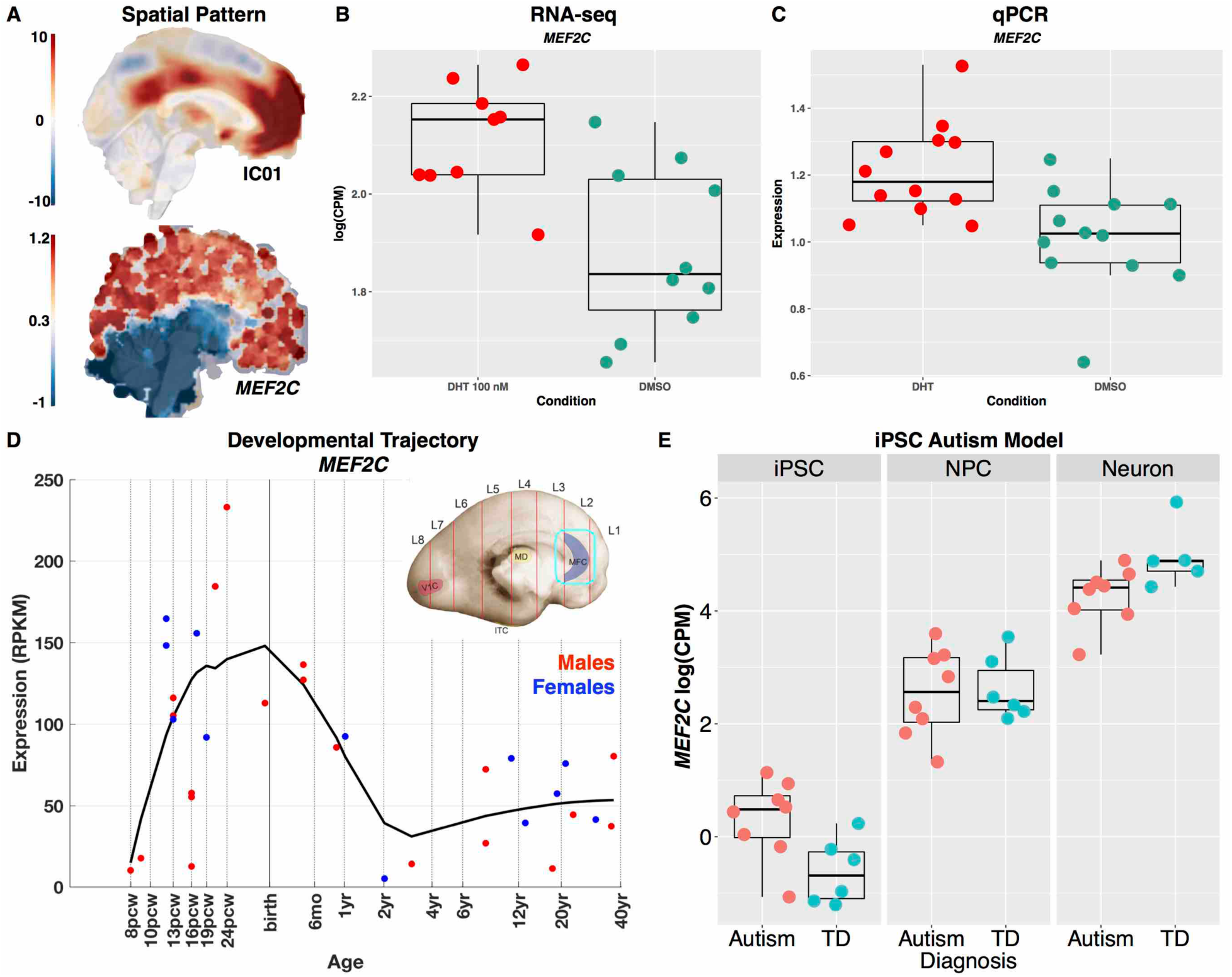
Panel A shows spatial pattern of MEF2C expression compared to spatial rsfMRI map for IC01. The whole-brain MEF2C expression maps shown is a composite map averaging across all donors. This composite map is shown for visualization purposes only. It is not meant to reflect directly the hierarchical statistical testing as implemented with Neurosynth Gene Expression Decoding. Panels B and C show expression of MEF2C across RNA-seq (B) and qPCR (C) experiments. Panel D shows the developmental trajectory of MEF2C expression in the Allen Institute BrainSpan atlas (blue, female; red male). Panel E shows MEF2C expression across induced pluripotent stem cells (iPSC), neural progenitor cells (NPC), and neurons from cases with autism or typically developing controls (TD).

## Discussion

In this study, we have shown that variation in testosterone during midgestational prenatal development can have long-lasting impact on how specific networks comprising the core of the social brain, the DMN, are organized. In line with hypotheses about sex-specific mechanisms, we have identified that the fetal programming impact of testosterone on later circuit-level organization of the DMN depends on whether an individual is male or female. These findings are compatible with other normative large-scale work on sex differences in DMN connectivity. Two large independent studies on adults utilizing the 1000 Functional Connectomes or UK Biobank datasets now replicably show that default mode connectivity is on-average stronger in females than in males^45, 46^. The directionality of these documented on-average DMN sex differences in functional connectivity are congruent with the directionality of the male-specific influence shown here - FT dampens later DMN connectivity, and thus could be one explanation behind the on-average lower DMN connectivity in males observed in other studies^45, 46^.

These findings are also particularly relevant for understanding how the prenatal environmental can shape early neurobiological mechanisms and heighten male-susceptibility for early-onset neurodevelopmental conditions. Autism is an early neurodevelopmental condition of particular relevance here, given the 3:1 sex ratio^12^ alongside other work implicating elevated prenatal steroidogenic activity^7, 17^. The hallmark features of autism are profound difficulties with early social-communication and social behavior, which has led to many investigations on the possible link with default mode network organization and function. The two core DMN subsystems identified in this work are known for the importance in both mentalizing and self-referential cognition and these domains along with the DMN circuitry underlying them are known to be atypical in males with autism^47-50^. The current results suggest that FT could act as a male-specific mechanism that dampens connectivity between core default mode subsystems linked to domains that are highly important for social-communication.

Showing that FT influences later functional organization within the human brain is the first step in understanding the developmental biology behind this effect. Imperative in working towards this translational goal, we need to understand how FT might have male-specific biological influence on developing neural circuits for the social brain, such as the DMN. Using a human neural stem cell model, we identified a subset of genes that are differentially expressed after androgen treatment. In an important link to autism, this set of genes is enriched in a number of high-impact genes known to be syndromic causes of autism and intellectual disability. This finding underscores a prior paper^25^ utilizing the same dataset that came across similar findings albeit with a different approach for differential expression analysis. However, this finding makes some specific inferences with regards to enrichments with the ‘syndromic’ category of variants (n = 102) labeled within SFARI Gene, whereas the prior paper looked for enrichment with any of n = 235 genes annotated within SFARI Gene.

In keeping with the relevant early phase of neurodevelopment for hNSCs, the DHT manipulation affected many genes involved in important neurodevelopmental processes including neurogenesis, cell differentiation, pattern specification and regionalization, and morphogenesis. These findings are also in line with previous work^25^ whereby independent experiments showed that DHT enhances cell proliferation and prevents cell death during neuronal differentiation under nutrient-deprived conditions. These processes are hypothesized to be some of the earliest key prenatal processes that are affected in autism and dysregulation of these processes can have multifinal outcomes and pleiotropic effects later in life including atypical circuit formation and function^51^. Androgen activity can be considered as an epigenetic influence over prenatal brain development given that such sex hormones work via androgen receptor signaling to directly affect transcription of many other genes. This work further supports the idea that androgens can exert epigenetic impact over prenatal neurobiological processes that are highly linked to male-biased conditions like autism.

Going beyond the list of DHT-dysregulated genes, we went further to pinpoint a subset of DHT-influenced genes that are also highly relevant specifically for the cortical midline default mode network subsystem that shows a male-specific influence of FT. This subset of DHT-dysregulated and cortical midline DMN-relevant genes was enriched for a variety of synaptic processes and potentially highlights important biological mechanisms in prenatal development that androgens act on. For example, the top enrichment in such genes was for excitatory postsynaptic potentials and includes genes for glutamate receptors such as *GRIK2*, *GRIA1*, and *GRIA2*, all of which are upregulated by DHT. Also included in this enrichment is *MEF2C*, which is upregulated by DHT, and is known to alter excitation/inhibition (E/I) imbalance^52, 53^. Also relevant to E/I imbalance is *NPTX2*, which has effects on GluA4-containing AMPA receptors^54^ and synaptogenesis^55^. DHT also upregulates sodium-gated ion channel genes (*SCN3A*, *SCN9A*), which have spatial similarity expression with cortical midline DMN and could also be relevant to E/I imbalance. All of these effects whereby DHT upregulates expression of such genes may point to the importance of E/I imbalance in androgen impact on early prenatal brain development. This is particularly relevant to autism given common theoretical views about E/I imbalance in autism^56^. Because these genes are highly expressed in spatial patterns resembling the DMN network we find associated with FT in rsfMRI, it may be that these are key mechanisms of prenatal androgen-impact on early developing social brain DMN circuit formation and maintenance.

While there are many genes from this work that could be focused on, we particularly highlight the role of *MEF2C*. Ample evidence supports the involvement of *MEF2C* as a candidate mechanisms involved in male-biased early developmental conditions such as autism and intellectual disability^4, 53, 57-60^. Developmentally, *MEF2C* is highly expressed during prenatal development (Figure 3D). Within prenatal development *MEF2C* has the potential to be a transcriptional regulator of a number of autism-associated genes - *MEF2C* binding motifs are enriched in upstream regions of genes within prenatal gene co-expression modules that harbor a number of autism-associated genes^4^. Congruent with the idea that *MEF2C* is important for many prenatal neurodevelopmental processes, other work has shown it is involved in neurogenesis, cell differentiation, maturation, and migration^61-63^ as well as having later roles to play in experience-dependent synaptic development and cell death^52, 53, 64, 65^. *MEF2C* is also a known downstream target of the androgen receptor (*AR*), thus making it highly susceptible to androgen-dependent transcriptional influence^43^. *MEF2C* is also one of the most important transcription factor genes influencing sex differences in human brain gene expression - *MEF2C* differentially targets genes as a function of sex, and the magnitude of this sex-differential targeting explains a large percentage of variance in sex differences in gene expression of its putative targets^44^. Our work further underscores *MEF2C*’s involvement in male-specific mechanisms behind atypical neurodevelopment, and particularly highlights how *MEF2C* could be one of many candidates behind fetal programming effects of androgens on developing social brain circuits such as the DMN. Compatible with effects on early neurogenesis, differentiation, and synaptic development, we find evidence that *MEF2C* expression is dysregulated in iPS cells and neurons in male cases with autism. Thus, important future work could examine *AR* targeting of *MEF2C* and its effect on early neurogenesis and differentiation processes as well as later synaptic processes at work in neurons.

In conclusion, we find that variation in FT during midgestational periods of prenatal development has a sex-specific impact on later human brain development. In particular, FT dampens functional connectivity in adolescence between social brain DMN subsystems in males, but has no effect on DMN functional connectivity in females. This sex-specific influence in early prenatal development was modeled in hNSCs to discover how androgens may act as a transcriptional influence on genes that are highly relevant in the adult brain for specific DMN-related circuitry. Here we discovered that DHT-dysregulated genes are highly enriched with a number of genes in the adult brain that are expressed spatially in a similar pattern to the DMN. These DHT-dysregulated and DMN-relevant genes and are involved in a variety of synaptic processes, congruent with the idea that FT may exert fetal programming influence on genes that play roles in biological processes that are integral for later circuit formation and maintenance. DHT also plays a prominent role in dysregulating many early neurodevelopmental processes such as neurogenesis, cell differentiation, patterning, and regionalization. This effect is compatible with the idea that androgens can affect genes that may have pleiotropic roles at early and later phases of brain development that link early cell proliferation and differentiation to later synaptic organization. *MEF2C* is one particular gene with such pleiotropic roles and is highly associated with male-biased conditions such as autism and intellectual disability. Congruent with predictions that *MEF2C* may be dysregulated in autism at both early and later phases of cell development, we find evidence suggesting that *MEF2C* is indeed dysregulated within iPS cells and neurons. This work highlights that prenatal androgens may have male-specific influence over early prenatal neurodevelopmental processes that can potentially manifest as long-lasting influence over social brain circuitry. These effects may help explain normative sex differences in brain and behavior as well as increased male-susceptibility to conditions such as autism.

## Acknowledgments

This work was supported by a Wellcome Trust (091774/Z/10/Z) project grant to SB-C and ETB and grants from the MRC, Autism Research Trust, and the Templeton World Charitable Foundation to SBC. This work is also supported by the National Institute for Health Research (NIHR) Collaboration for Leadership in Applied Health Research and Care (CLAHRC) East of England at Cambridgeshire and Peterborough NHS Foundation Trust. The views expressed are those of the authors and not necessarily those of the NHS, the NIHR, or the Department of Health. MVL and BA were supported by the Wellcome Trust. MVL was also supported by the British Academy, Jesus College Cambridge, and the European Research Council (ERC; 755816) during this period of work. AP and J-LM were supported by a grant from the Agence Nationale de la Recherche (HARTAGeNe). AQ also received additional support from a fellowship from the Fondation pour la Recherche Médicale. TP was supported by a Brain & Behavior Research Foundation (NARSAD). M-CL and AR received support from the William Binks Autism Neuroscience Fellowship at the University of Cambridge. M-CL was supported by the O’Brien Scholars Program within the Child and Youth Mental Health Collaborative at the Centre for Addiction and Mental Health and the Hospital for Sick Children, Toronto, and the Slifka-Ritvo Award for Innovation in Autism Research by the Alan B. Slifka Foundation and the International Society for Autism Research. PK was supported by the National Institutes of Health–Cambridge Scholars Program. We would also like to thank Eric Courchesne for valuable discussion and comments.

## Author Contributions

MVL, BA, PK, ETB, and SBC designed the neuroimaging experiment for this work. MVL, BA, RJH, JW, NM, and ANVR implemented the neuroimaging experiment. AP, J-LM, AQ, and JC designed and implemented all experiments on hNSCs. MVL conceived and implemented all analyses. TP contributed data for autism iPSC experiment, bioinformatic analyses, and interpretation of results. All authors were involved in drafting the manuscript.

## Supplementary Results and Figures

Quartier and colleagues recently reported DHT dysregulates genes were enriched for autism-associated genes^25^. The DE analysis in this paper utilizes the same RNA-seq dataset, but performs the DE analysis differently than that of Quartier and colleagues. Therefore, we reexamined this question with the current study’s DE lists. Upon examining SFARI Gene Scoring categories for overlap with autism-associated genes, we find that a number of genes in the ‘Syndromic’ category are enriched in genes that are influenced by DHT manipulation (OR = 2.16, p = 0.02). These genes include *AHI1*, *ASXL3*, *CHD2*, *DHCR7*, *HCN1*, *MEF2C*, *PAX6*, *PRODH*, *PTEN*, and *SCN1A*. *ASXL3* and *PTEN* are also classified in the high-confidence category. No other gene scoring categories were enriched. However, there were many other notable autism-associated genes influenced by DHT, including *NLGN4X*, *NRXN3*, *FOXP1*, and *SCN9A* (Supplementary Figure 1C). Thus, this analysis highlights a largely-overlapping finding with the prior paper by Quartier et al.,^25^ showing that many syndromic causes of autism are enriched within the DHT-dysregulated gene set.

**Supplementary Figure 1:**
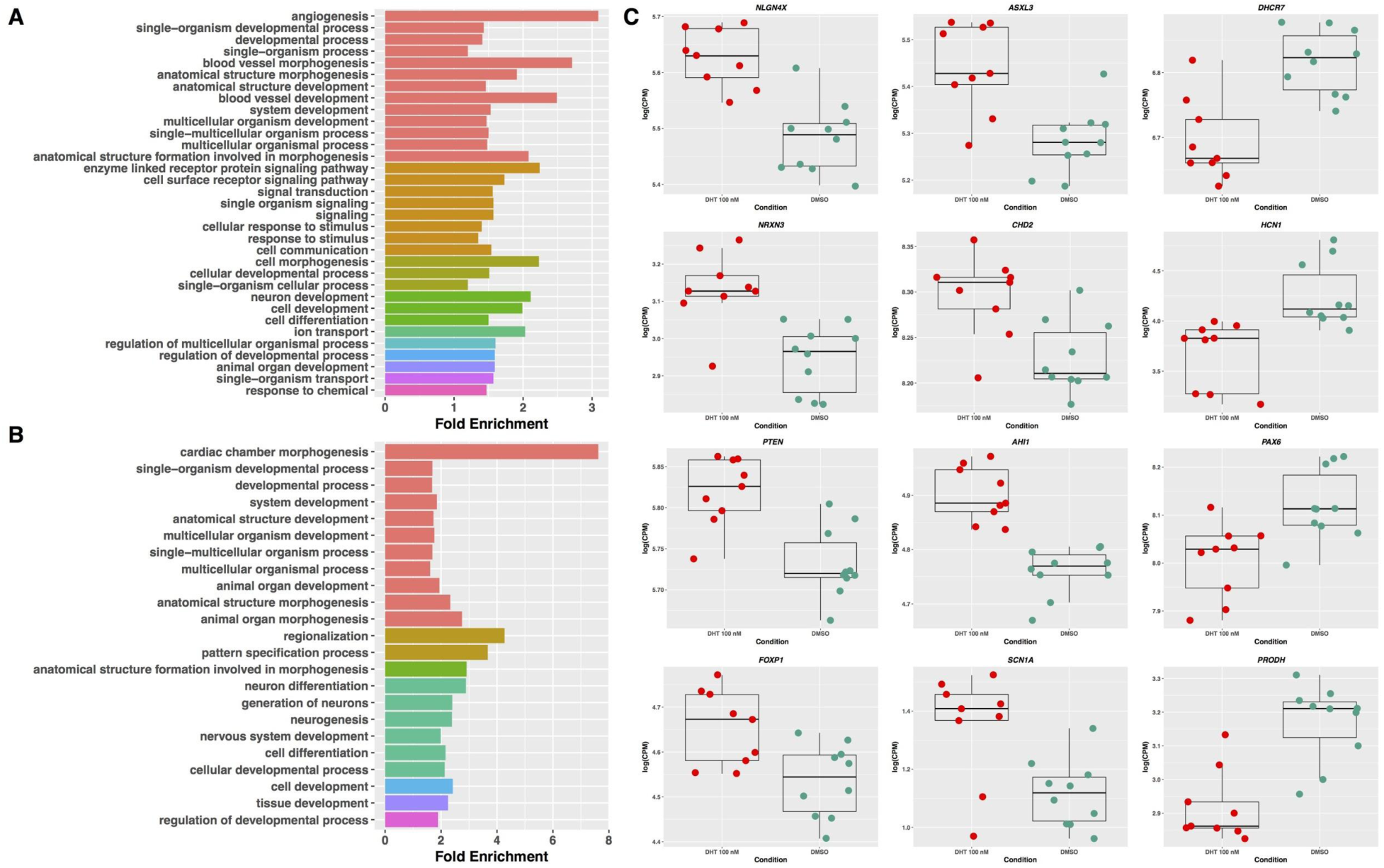
Biological process enrichment of DHT-upregulated (A) and downregulated (B) genes sets and plots of specific autism-associated genes (C).

